# *dar*: A Consensus-Based Framework for Differential Abundance Testing in Microbiome Data

**DOI:** 10.1101/2025.02.07.637000

**Authors:** Judit Farré-Badia, Marc Noguera-Julian, Roger Paredes, Francesc Català-Moll

## Abstract

**Summary:** The *dar* R package streamlines differential abundance (DA) testing in microbiome research by integrating state-of-the-art DA methods—such as DESeq2, ALDEx2, ANCOM-BC, and MetagenomeSeq—within a customizable consensus-based framework, thereby enhancing the robustness and reproducibility of DA results. Leveraging microbiome data in *phyloseq* or *TreeSummarizedExperiment* formats, *dar* organizes analysis steps in a modular *recipe* object, enabling users to easily incorporate preprocessing tasks such as taxonomic filtering, rarefaction, and subsetting, alongside multiple DA analysis methods. Dedicated visualization tools facilitate the definition of a consensus by illustrating the overlap among methods, empowering users to refine analysis strategies and display final results. Reproducibility is supported through functions that export and import entire analysis workflows, making *dar* a comprehensive solution for addressing the complex, high-dimensional nature of microbiome data.

**Availability and implementation:** *dar* is an R package available from Bioconductor ≥ 3.19 (https://www.bioconductor.org/packages/dar) for R ≥ 4.4. The software is distributed under the MIT License and includes example datasets.

**Supplementary information:** Additional documentation is available at https://microbialgenomics-irsicaixaorg.github.io/dar

## 1. INTRODUCTION

Microbiome data analysis is a rapidly growing field with numerous applications in the study of the complex interactions between microorganisms and their hosts, with important implications for health and disease (Gilbert *et al*., 2018; Cho and Blaser, 2012). A key aspect of microbiome analysis is differential abundance (DA) testing, which identifies differences in taxa abundances across sample groups or conditions (Paulson *et al*., 2013; The Human Microbiome Project Consortium, 2012). DA analysis is a valuable tool for uncovering the mechanisms underlying observed differences in health outcomes, while also enabling the identification of potential biomarkers and therapeutic targets. For instance, variations in the abundance of specific bacterial taxa have been linked to conditions such as obesity, type II diabetes, and inflammatory bowel disease (Ley, 2010; Qin *et al*., 2012; Franzosa *et al*., 2018)..

However, DA analysis in microbiome research is inherently challenging due to high data sparsity, variability, and the compositional nature of microbiome data. Such datasets are often high-dimensional, zero-inflated and characterized by low counts across many taxa (Paulson *et al*., 2013; McMurdie and Holmes, 2014; Morton *et al*., 2019; Weiss *et al*., 2017). Moreover, relative proportions of taxa within a sample can be more biologically meaningful than absolute abundances, meaning conventional statistical approaches may be insufficient (Lovell *et al*., 2015; Aitchison, 1982; Gloor *et al*., 2016).

Although many DA methods —such as ALDEx2, ANCOM-BC, and MetagenomeSeq—have emerged, no single approach consistently outperforms the others across all data types and experimental contexts. Consequently, relying on one DA method can be problematic, and recent benchmark studies recommend using multiple methods to obtain reliable results (Nearing *et al*., 2022; Yang and Chen, 2022). To facilitate the adoption of consensus-based strategies, we developed *dar*, an R package that unifies multiple DA tools under a customizable consensus framework, thereby enabling robust and reproducible DA testing. Additionally, *dar* allows users to export and import complete analysis workflows, further promoting reproducibility in microbiome data analyses.

## 2. IMPLEMENTATION

### 2.1 Recipe Object Structure and Data Organization

In *dar*, the analysis workflow is managed via a recipe object—an S4 class instance—that sequences both preprocessing and DA tasks as modular “steps” (see Fig. 1A). Each step corresponds to a specific data transformation (e.g., filtering, rarefaction, or DA analysis) and is fully customizable. This modularity provides users with the flexibility to reorder or omit steps based on the characteristics of their dataset.

**Figure 1.**
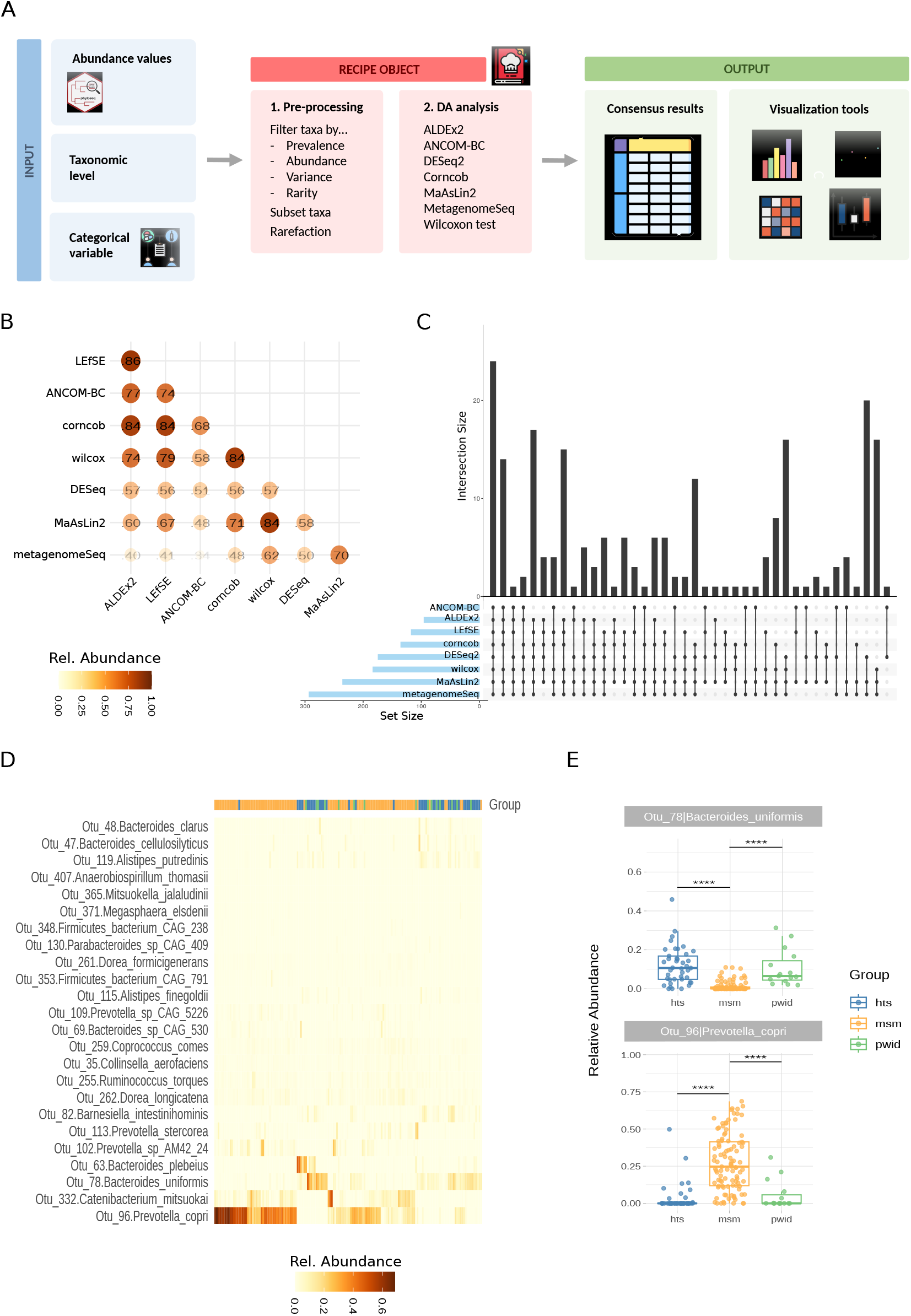
Overview of the *dar* Workflow and Case Study Results. **(A)** Schematic diagram illustrating the modular recipe object structure in dar, which sequentially organizes data preprocessing and differential abundance (DA) analysis steps. **(B)** Correlation heatmap comparing the outcomes of multiple DA methods, highlighting the consistency and divergence among methods. **(C)** Intersection diagram showing the overlap of significant taxa identified by various DA methods under different consensus thresholds, demonstrating how adjusting the cutoff impacts the number of shared taxa. **(D)** Heatmap of the final consensus taxa, confirming differential abundance patterns between sample groups (e.g., higher Prevotella levels in MSM vs. increased Bacteroides in HTS). **(E)** Boxplots displaying the abundance distribution of selected taxa across sample groups, providing a detailed view of the differential abundance validated by the consensus approach.

The recipe object is initialized with a phyloseq or TreeSummarizedExperiment object that contains the primary microbiome data (counts, taxonomic classifications, and sample metadata). Users specify the main categorical variable (e.g., health status or environmental factor) to define DA contrasts and the taxonomic level (e.g., species or genus) for the analysis. Subsequent steps—either for data preprocessing or DA method execution—are then appended to the recipe object.

### 2.2 Data Preprocessing and Differential Abundance Analysis

Upon initialization, a preliminary data quality report is generated to help users determine appropriate preprocessing measures. The recipe object supports multiple filtering procedures, including taxonomic filtering to retain taxa of interest, abundance filtering to remove taxa that are too rare or have low prevalence, and rarefaction to equalize library sizes across samples.

Users may then append any combination of the DA methods supported by *dar*—for example, DESeq2, ALDEx2, ANCOM-BC, MetagenomeSeq, corncob, LEfSe, MaAsLin2, and the Wilcoxon test—and customize their respective parameters. By invoking the prep function, *dar* sequentially (or in parallel) executes all preprocessing steps and applies the selected DA methods to the updated dataset.

### 2.3 Consensus Building for Robust DA Results

A central feature of *dar* is its customizable majority-vote consensus approach, which integrates outcomes from multiple DA methods to reduce method-specific biases. Users can define consensus thresholds—for instance, requiring a taxon to be identified as significant by a minimum number of methods—to improve the reliability of the findings.

Visualization plays a crucial role in interpreting and refining the consensus strategy. The *dar* package provides graphical methods to explore how different methods converge or diverge in their calls, including intersection diagrams to illustrate shared significant taxa, correlation heatmaps to reveal similarities among methods, and abundance plots for summarizing selected features.

After choosing a consensus strategy, the *bake* function applies the user-defined criteria, and the final results are retrieved with the *cool* function. Users can also export or import the entire recipe object—encompassing all parameters—thereby promoting reproducibility and transparency in microbiome analyses.

## 3. CASE STUDY

To demonstrate the capabilities of *dar*, we performed a case study analyzing gut microbiome differences based on sexual orientation within a human cohort (Noguera-Julian *et al*., 2016). This dataset exemplifies common challenges in microbiome research, such as zero-inflation and high inter-individual variability.

First, the data were loaded into a *recipe* object and processed at the species level. Samples were categorized into three groups: men who have sex with men (MSM), non-MSM (HTS), and people who inject drugs (PWID). Taxonomic filtering was applied using a prevalence threshold (≥1% in at least one sample), and the analysis was restricted to Bacteria and Archaea to ensure a high-quality dataset.

Subsequently, we appended eight DA methods—DESeq2, ALDEx2, ANCOM-BC, LEfSe, corncob, MaAsLin2, the Wilcoxon test, and MetagenomeSeq—to the recipe object. After executing the *prep* function, we assessed the reliability of the results using *dar*’s visualization toolkit. The correlation heatmap (Fig. 1B) revealed that most methods produced consistent outcomes, although DESeq2 and MetagenomeSeq exhibited relatively lower correlations with the other methods, suggesting potential divergences in modeling assumptions. Initially, 24 taxa were consistently identified across all methods. By adjusting the consensus threshold to require significance in at least six methods, the number of shared taxa increased to 58 (Fig. 1C), illustrating the impact of threshold settings on the results.

Finally, after defining a consensus strategy that retained only taxa identified as differentially abundant by all tested methods, we visualized the abundance of the resulting 24 taxa using a heatmap (Fig. 1D). This confirmed previous findings: MSM individuals typically harbored higher levels of *Prevotella*, while HTS subjects exhibited greater *Bacteroides* abundance. In addition to heatmaps, *dar* supports boxplots for detailed inspection of each taxon’s abundance distribution (Fig. 1E).

## 4. DISCUSSION

The *dar* package offers a comprehensive solution for differential abundance testing in microbiome research. By integrating multiple DA methods within a single, customizable framework, *dar* overcomes the limitations inherent in single-method approaches when analyzing high-dimensional, compositional data (Aitchison, 1982; Nearing *et al*., 2022; Yang and Chen, 2022). Its consensus-driven approach aligns with current recommendations for employing multiple methods to achieve more accurate and reliable results, while also enabling straightforward method comparison.

The modular design of *dar* allows users to tailor analysis workflows to their specific needs. Its seamless integration with the R/Bioconductor and tidyverse ecosystems further enhances reproducibility and accessibility. Nevertheless, some limitations remain. For example, applying multiple DA methods concurrently may increase computational demands. Future work should focus on optimizing computational efficiency and exploring alternative consensus criteria.

## DATA AVAILABILITY

All data presented in this manuscript have been previously published (Noguera-Julian *et al*., 2016) and are accessible either via the original publication or as internal data within the package.

## CODE AVAILABILITY

The complete code used to generate Figures 1B, 1C, 1D, and 1E is available at dar’s GitHub repository. https://microbialgenomics-irsicaixaorg.github.io/dar/vignettes/bioinformatics_vignette.qmd

## ACKNOWLEDGEMENTS

We express our heartfelt thanks to the Bioconductor community for its invaluable support and resources. We are especially grateful to those dedicated to developing bioinformatics tools specifically for microbiome research, whose contributions have significantly advanced this field.

## AUTHOR CONTRIBUTION STATEMENT

JFB: Software, Formal analysis, Writing – Original Draft, Writing – other versions, Visualization.

MNJ: Conceptualization

RP: Supervision, Project administration, Funding acquisition.

FCM: Conceptualization, Methodology, Software, Formal analysis, Investigation, Resources, Writing – Original Draft, Writing – other versions, Visualization.

